# Using vocational education to provide development solutions in the Pacific: An emphasis on climate change and health

**DOI:** 10.1101/350009

**Authors:** Peni Hausia Havea, Amelia Siga, Titilia Rabuatoka, Apenisa Tagivetaua Tamani, Priya Devi, Ruci Senikula, Sarah L Hemstock, Helene Jacot Des Combes

## Abstract

This article reports on the results of the EU PacTVET project, which explored the use of Technical Vocational Education and Training (TVET) to provide a better understanding on the development solution for the impact of climate change on human health in the region. It describes the findings of a 2014-2018 project on the use of vocational education to provide development solutions in the Pacific with an emphasis on climate change and health. An exploratory design was used to investigate how vocational education developed solutions for climate change and health in the 15 Pacific – African Caribbean and Pacific (P-ACP) countries: Cook Islands, Federated States of Micronesia (FSM), Fiji, Kiribati, Nauru, Niue, Palau, Papua New Guinea (PNG), Republic of Marshall Islands (RMI), Samoa, Solomon Islands, Timor-Leste, Tonga, Tuvalu and Vanuatu. Information collected via personal communication with relevant stakeholders, qualitative interviews, documents review, and survey (n=48) of youths and young women in Fiji. Data analysis was performed using thematic analytical strategy and frequency analysis. The study found that vocational education plays a significant role in building the capacity of people to become more sustainable and resilient in their life now and in the future. Also, getting an accredited qualification on health resilience and/or job in the health sector may help them to respond effectively and efficiently in the event of climate change and/or disasters caused by natural hazards. The same factors were explored quantitatively using descriptive analytical strategy, and concluded TVET education, to have a positive influence on climate change and health. As a result, vocational education could provide development solutions for health adaptation in the Pacific. These results indicate global actions for vocational education, that would perfect the course of resilience for these 15 P-ACP in the Pacific and alike in the U.S.

## Introduction

Global climate change and disasters caused by natural hazards are known to affect many sectors worldwide including the Pacific [1–11]. In the Pacific, these sectors may include but not limited to agriculture, coastal management, energy and infrastructure, education, fishery, forestry, health, tourism, and water resources [11–22]. As a result of these phenomenal impacts on all levels of society for the people of the Pacific, all Pacific governments, Non-Government Organisations (NGOs), regional and international organisations were then mandated to respond to the regional countries, in order to enhance their sustainable development solutions to build more resilient Pacific Islanders, by 2030 and beyond.

Such a confluent for actions is significant to the livelihoods, health and well-being of the people in the region, leaders in the Pacific then envisioned the birth of the EU PacTVET project in August 2014 with an overall budget of EUR 6.1 million, as part of its worldwide contribution to adapting to climate change (CCA) and hazards (DRR) and Sustainable Energy (SE) development in the region. The program was specifically designed to enhance sustainable livelihoods, thus strengthening countries’ capabilities to adapt to the adverse effects of climate change as well as enhancing their energy security at all levels in 15 Pacific Island Countries (PICs): Cook Islands, Federated States of Micronesia (FSM), Fiji, Kiribati, Nauru, Niue, Palau, Papua New Guinea (PNG), Republic of the Marshall Islands (RMI), Samoa, Solomon Islands, Timor-Leste, Tonga, Tuvalu and Vanuatu (Fig 1).

**Fig 1.**
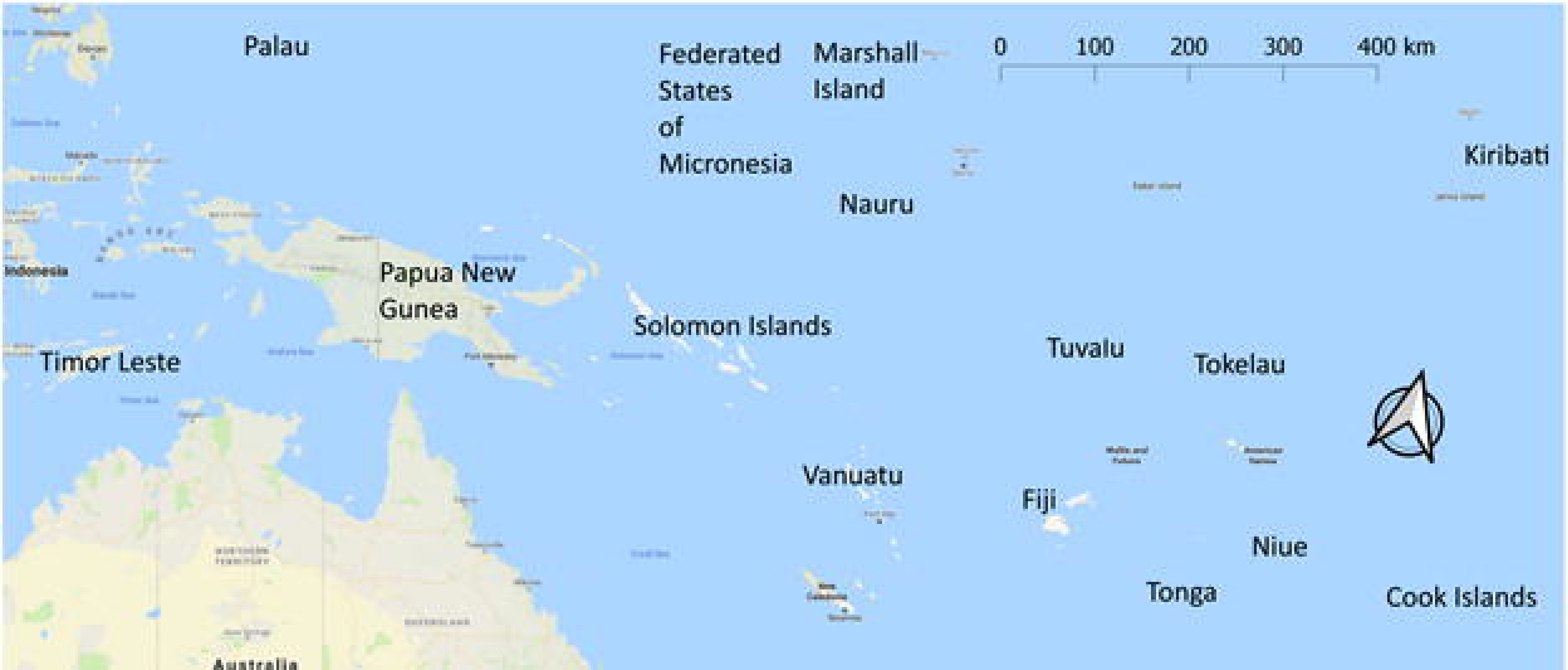
Map of the 15 Pacific Island Countries who are participated in the EU PacTVET Project.

The 10th European Development Fund European Union Pacific Technical and Vocational Education and Training on Sustainable Energy and Climate Change Adaptation (European Union PacTVET) project is component three within the broader regional Adapting to Climate Change and Sustainable Energy (ACSE) programme. Both the EU and GIZ are EU PacTVET project implementing partners. The project builds on the recognition that energy security and climate change are major issues that are currently hindering the social, environmental and economic development of Pacific – African Caribbean and Pacific (P-ACP) countries [23]. It is also the first programme in the region to combine both resilience (CCA & DRR) and sustainable energy in a single project.

In the Pacific, to date only the World Health Organisation [14, 24] and the Government of Fiji through the Ministry of Health and Medical Services [25] have had programmes emphasising climate change and health. The WHO report on “Human Health and Climate Change in Pacific Island Countries” in 2015 focused on 13 PICs: Cook Islands, FSM, Fiji, Kiribati, RMI, Nauru, Niue, Palau, Samoa, Solomon Islands, Tonga, Tuvalu, Vanuatu. The assessment was on vulnerabilities to the impacts of climate change on health and adaptation strategies. Since Fiji was part of the WHO project in 2016, this led to the Government of Fiji developing “Climate Change and Health Strategic Action Plan 2016-2020” for the Climate Change Unit of the Ministry of Health and Medical Services (MoHMS). The other PICs are still in the process of developing, their own national health adaptation plans. For example, The Queen Salote Institute of Nursing and Allied Health in Tonga and the School of Nursing in Fiji are planning to integrate climate change and health into their curriculum in the future.

Since most of the climate change and health programmes are only available at the graduate level (e.g. master and Ph.D. etc.) at universities like the University of the South Pacific (USP) and Fiji National University (FNU), the deficiency of literature on this topic indicates that it is currently under-researched. At the outset of the EU PacTVET project, there were no formal vocational qualifications in this area, with ten of the fifteen countries the project is working in having no functional national vocational educational quality assurance systems [26]. Significantly, the EU PacTVET project embarked not only to develop solutions for climate change and health via vocational education by developing accredited qualifications on Health and Resilience (CCA & DRR) from certificate level 1 to certificate level 4, it is also working with the Educational Quality Assessment Programme of the Pacific Community on regional accreditation on institutional verification so that the qualifications can, in theory, be delivered at any TVET institution in the region.

According to the Pacific Association of Technical and Vocational Education and Training (PATVET) [27], 12 countries have vocational institutions: Cook Islands, Fiji, Kiribati, Nauru, Niue, PNG, RMI, Samoa, Solomon Islands, Tonga, Tuvalu and Vanuatu – and most of these countries are USP members countries, where they also host the Pacific Technical And Further Education (TAFE) course on Resilience programme in their regional campuses. Additionally, since Fiji has more than 100 vocational schools/institutions and PNG more than 130, using the EU PacTVET project as a guide via its 15 PICs, it is expected that the vocational education in these countries, as well as USP, could regionalize and/or revolutionalise the development solutions for climate change and health in the Pacific, especially at the grassroots level such as primary, secondary and TVET education. The by-product of this initiative is that it may not only help to fill in this gap in knowledge and needs for research in the region, but also help to build Pacific Island communities that are more sustainable and resilient now and in the future.

This may lead not only to perfect the course of climate change and health adaptation but also contribute to the achievement of the Sustainable Development Goals (SDGs), which has health as goal number 3 [28], the targets and objectives of the United Nations Framework Convention on Climate Change (UNFCCC), Sendai Framework for Disaster Risk Reduction 2015-2030 and Framework for Resilient Development in the Pacific, by 2030 and beyond.

## Methods

### Methodology

A mixed method approach, named explanatory design [29], was used to gather all the quantitative aspects of the EU PacTVET project. This quantitative data was collected from the 15 Pacific – African Caribbean and Pacific (P-ACP) countries in the region where the project was implemented. The quantitative data for the project was mainly from surveys, registration of students in TVET institutions and health expenditure. The qualitative aspect of the project was mainly information that was collected from personal communications, interviews and project documents. The study approach was called explanatory [30] because this paper has relied heavily on the quantitative aspect of the project.

### Data Analytical Strategy

The data analysis used an explanatory design model [31]. For the quantitative data, the analysis used frequency. The results of this analysis were then explored qualitatively using thematic analytical strategy. As a result, the quantitative results were explored qualitatively and vice versa in order to provide a better understanding of how to use TVET education to develop solutions for health and climate change. The data analysis was performed using SPSS and r studio.

## Results

Based on the results of this project, the study found four development solutions that TVET education has used to help people in the Pacific improve their capacity to address and to prevent some of the worst impacts of climate change on health and ultimately their well-being.

### 1) Using the TVET Education Model to Address Development Solution on Climate Change and Health

From studies across the 15 P-ACP countries, a key barrier to the development of the first qualification on resilience and health is because of no formal vocational sector qualifications in CCA-DRR (Resilience) prior to the existence of the EU PacTVET project. This has led to the development of the EU PacTVET model to address this limitation. This has been done by integrating health and climate change into certificate levels 1-4 for TVET education in resilience.

Using the EU PacTVET project’s state-of-the-art ideology, a climate change landscaping model has factored health into this conceptual framework. This state-of-the-art idea was used to develop a framework on climate change and health for the 15 participating countries, that not only enhance the development solutions of their people in their respective countries but equip them to achieve sustainable and resilient health and happy well-being now and in the future (Fig 2).

**Fig 2.**
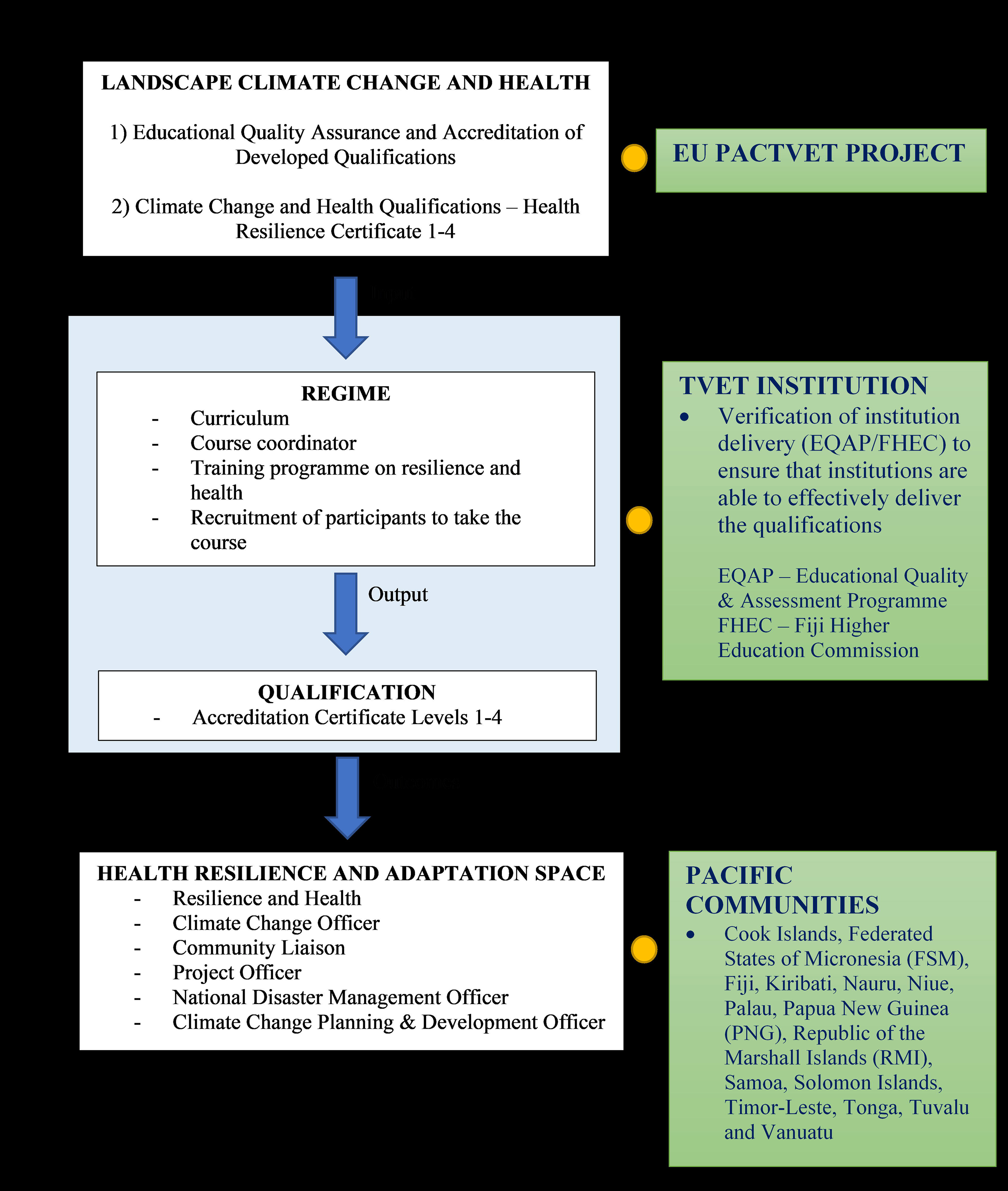
EU PacTEVT Model for Accredited Qualification on Certificate Level 1-4 on Resilience and Health.

In this model, there were four main steps: 1) identification of climate change landscaping (development solution – climate change and health); 2) climate change regime; 3) qualification on resilience in health; 4) and achievement of healthy living, sustainable and resilient Pacific Islanders (adaptation space), by 2030 and beyond.

#### 1) <u>Landscaping Climate Change and Health into Certificate Levels 1 to 4 in TVET Education</u>

In this context, landscaping climate change and health may mean using the insight of landscape dynamics to unfold challenges that are difficult to embrace empirically by formalizing qualifications in the forms of accreditation and quality control for climate change and health in certificate levels 1 to 4. As a result, it is the science of studying to better understand the relationship between climate change and health, thus developing a solution that may contribute to solving health problems in Pacific communities. This landscaping of climate change and health for the region may represent the EU PacTVET project, since, it is the project, that developed these qualifications to be recognized nationally and regionally by the national and regional qualifications frameworks that are already in place.

#### 2) <u>Climate Change Regime</u>

When the process of landscaping the qualification for climate change and health was completed by the project, the next phase was to input it into the climate change regime. For the purpose of this project, this was inputted in the form of niche or in the form of innovation to endogenous factors such as the development of the curriculum for certificate levels 1 to 4 on climate change and health and exogenous factors, such as the recruitment of a qualified course coordinator to coordinate and facilitate this programme. In this project, the regime may represent an organisation or TVET institution.

#### 3) <u>Climate Change and Health Qualification</u>

Within the TVET institution or an organisation, the next phase of the model is the delivery of the qualifications to the learners or to the participants as an output based on the regime. These institutions are verified by regional accreditation agencies such as Education Quality & Assessment Programme (EQAP) and/or Fiji Higher Education Commission (FHEC) to deliver accredited-based qualifications. These accredited qualifications on climate change and health are available for delivery by the 15 P-ACP countries that were part of this project. Some of the countries started with this delivery in 2017. The qualifications were accredited because they met the Pacific Qualification Framework (PQF) standards as well as international standards like the Australian Qualification Framework (AQF) on certificate levels 1 to 4. Nationalization of qualifications for primary and secondary education or certificate levels 1 and 2 on health and climate change is significant because it reveals much more of an individual level of competency to pursue and advance sustainability and resilience in life and/or the development solutions to their own problems than just their academic prowess.

#### 4) <u>Adaptation Space</u>

When the above-mentioned steps were completed, the final step is the outcome of delivering these qualifications. The outcome here is that the participants who represent the adaptation space will then be able to contribute, to providing solutions to their problems, be it at the community level, church, work environment or at the national level. Health resilience and adaptation space represents any solutions that people, who successfully completed the qualification(s) developed by the project, may have or use to protect themselves and others from the negative effects of climate change and hazards and then use their competency to shape their roles in climate and health adaptation for the benefits of all people in the Pacific.

Space may mean an adaptation space has been created for people in the Pacific to adapt to the effects of climate change in variable ways, to live a healthy life, thus achieving a resilient Pacific community by 2030 and beyond. Adaptation space may include time, space, money, being innovative and efficient in how they adapt inter alia, in order to achieve their goal in life. In summary, this is how the EU PacTVET project addresses the development solution for climate change and health in the form of formalizing the qualification for certificate levels 1-4 in resilience and health. For example, by the end of the training on resilience certificate level 4 at the USP Pacific TAFE, the learner will be competent to work as a:

- Climate Change Officer;
- Community Liaison;
- Project Officer;
- National Disaster Management Officer;
- Climate Change Planning & Development Officer;

amongst others, thus contributing to improving the adaptation space in the region. Adaptation space is the last phase of this climate change landscaping model because the adaptation is open to all to pursue the best option available to them.

### 2) Helping Pacific Countries to Incorporate Health and Climate Change in all Subjects in the Primary and Secondary Education Curriculum

The EU PacTVET project has been partnering with FHEC as one of its pilot areas for delivering the qualifications. EU PacTVET relied on the FHEC policies and procedures to initially develop and accredit the Regional Qualifications in Resilience (CCA and DRR). The project has also contributed to the enhancements of the primary and secondary education systems in the region in that some of the countries are planning to integrate climate change in all subjects for their primary and secondary education curriculum. For EU PacTVET, this has been achieved by partnering with GIZs Coping with Climate Change in the Pacific Islands Region (CCCPIR) project. If this initiative is successfully implemented, the Pacific region will be the first region in the world to streamline and/or teach climate change across all subjects at the primary and secondary levels.

For example, in Fiji, the Ministry of Education is planning to streamline climate change in all subjects in primary and secondary education including industrial arts, home economics, agricultural science, office technology, computer education (Fiji’s Ministry of Education 2018, personal communication, 21 March) accounting, social sciences, sciences, health study, mathematics, inter alia (A Tamani 2018, personal communication, 8 March). It is expected, by nationalizing and incorporating climate change and health into the school curriculum, it may help to build capacity on climate resilience and adaptation on health for the 15 P-ACP countries, thus guaranteeing a Pacific community that will be more resilient and sustainable by 2030 and beyond. In doing so, coding (to assign a code (numeric value or theme) for classification and/or identification of something (e.g. converting a message or text) in resilience in climate change should also be integrated into the school curriculum. Coding is important because it closes the gap between climate change and Information and Technology (IT).

### 3) Using TVET Education to Address Gender Towards Better Health and Resilience

The project also addressed gender equality – for women, girls and vulnerable groups (e.g. people with disability, etc.) – with the aim of developing a solution to improve skills for resilience and sustainable energy using TVET. From what is known in this project, although male learners dominated the TVET sector, the proportion of female learners is increasing for activities directly using the same learning medium to improve their well-being and to have a sustainable and resilient life now, and in the future (Fig 3).

**Fig 3.**
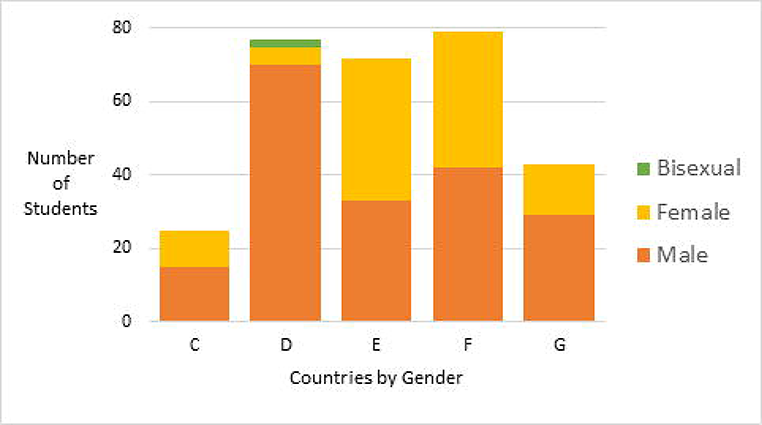
Number of students enrolled in TVET institution for 5 countries from years 7-13.

For example, a study by the project on 48 youths from communities in Fiji has shown that these young people chose climate change and health as their study priority if such a course or programme were to be offered in their schools or TVET institutions (Fig 4).

**Fig 4.**
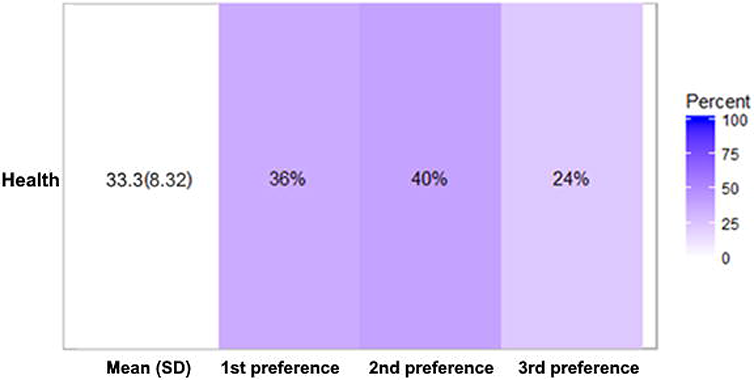
Percentage of youths and young women who preferred to study climate change and health, Fiji.

Although the project has allocated funding to address the needs of women, girls and vulnerable groups to recognize their rights to education and equal opportunity to climate resilience and adaptation on health, the Pacific community needs to highlight gender rights in its human rights convention, in order to widen the Pacific democratic space and work towards achieving SDG goal 5 on gender equality [32–35]. As a result, it will give not only respect for human rights – to perform unwittingly – but also a right to TVET education, to contribute significantly to gender equality, in achieving better health and resilience for the people of the Pacific.

### 4) By Helping Pacific People to Improve their Health Adaptation Strategies in their Communities – to Restore Better Health and Well-being

By giving formal education on climate change and health to the participating countries, it is expected that this will help the people of the Pacific to improve their health adaptation strategies in their communities to restore better health and well-being. This can be done in two ways. First, is to work together with the PATVET members countries since there are at least 12 PICs in this programme. They can do this through their Curriculum Development Units, which are mandated under the Ministry of Education – to improve basic knowledge and understanding for children and young people. Second, is to leverage the level of awareness for the general population. This can be done by integrating climate change and hazards into the census for these PATVET members countries. These are very important assets for the people of the Pacific.

People who will be learning climate change and health at TVET institution to become climate change officers, community liaison officers, project officers, national disaster management advisors and/or climate change planning and development officers, will most likely be more competent to live a sustainable and resilient life than those with an ad hoc adaptation skill. This can be applied to those who have participated in any projects regarding climate change impacts and adaptation strategies to coastal communities, since people will tend to have better solutions, and know how to deal with their problems better than those who do not learn about impacts of climate change on health and health resilience and adaptation [12, 13, 23, 36–40].

As one of the participants from ‘Ahau in Tongatapu, Tonga indicated when he was asked to reflect as to what extent a better understanding of the impact of climate change on livelihoods, health and well-being helps him as an individual, to prevent the risk of running into the same problems now and in the future stated: “This is a very important question. There was another important question, that pops up in the survey about recommending a policy proposal to prioritize a national survey about the impact of climate change on livelihoods, health and wellbeing in Tonga. To me personally, I think that is very important because we need to pass on this knowledge and learn how to deal with these problems like our elders. That’s how we are going to minimise these problems. So, that’s how I am going to help the next generations to come. Most importantly, this project really gives me some sort of insights into how to look after my families as well as my friends. As we speak earlier in our discussion today, we have been faced with this problem for so many years, regarding our homes and in this village. But for me, it gives a lot of better understanding how to deal with this problem better for now and in the future.”

## Discussion

### Training programme to focus on training experts in health adaptation

From what is known in the Pacific region, no TVET institution has used their educational programme to develop solutions for climate change and health. But this is an important solution for the landscaping of climate change because vocational education can train people to become experts in health resilience (CCA & DRR) at a lower level and their exposure to this level of training at a young age, will help them to build a resilient Pacific community. This is reiterated by the FRDP which identifies training and education, among other forms of human resources development, as vital to developing resilient communities whose members can actively engage in risk reduction activities and protect the interests of their most vulnerable population [41].

More importantly, this is the hallmark of building resilience in the Pacific – to empower those to whom resilience matters, as Paul Farmer called it the pathology of power [42]. It is poor and disempowered in the society, those who do not work, lack education and the inputs and systems of support to protect themselves from the negative impacts of climate change and hazards on their health in the region.

Seeing climate change and hazards in terms of frequency such as too many people were affected in the Pacific is too limited an approach to the problems. There is also a need to look at the social and economic conditions, cultural practices, religion and system of learning where people can be trained on resilience to reach where they live and work or to safeguard living “within their comfort zone”. To achieve this, training people on resilience in health through vocational education should not only welcome all people at all levels of society but with a purpose to determine the climate change determinant of health and to support the scaling up of health security.

This intervention will break the health impacts cycle by empowering people’s solution to adaptation and improving the footprint of sustainable living of those who will most likely be affected in the sense of changing the reality when the impacts or when the problems are happening. For example, by attending this training programme, people will be more competent about how to live their life in their communities as well as more confident about how to save themselves and others when there is a significant sea level rise or cyclones.

### Provide accredited qualifications on resilience in health

As the Pacific is constantly affected by climate change and natural disasters caused by natural hazards, vocational education in the region could come up with a common denominator between TVET, climate change and health which is to provide accredited qualifications on certificate levels 1 to 4 in Resilience (CCA & DRR). Significantly, this can be used by vocational education as part of their climate change regime to 2030 and beyond by creating more jobs that are secure and improving the conditions of employment that will lead to improvement in health adaptation and resilience. The EU PacTVET project can provide this competency-based education model to help the 15 P-ACP countries to meet not only education-related goals but to also address the shortages of skilled workers that are emerging to improve both health economics and resilience.

This is why the project made a significant partnership with the global leader in health such as WHO regional office in Suva, and regional health leader such as the Ministry of Health and Medical Services Department of Climate Change, School of Nursing and the School of Public Health at the Fiji National University in Fiji in developing these resilience and health certificates, to ensure that these sustainable goals will be met by 2020 and beyond. Most importantly, the consultant for the development of these resources were purely Pacific Islanders people, who are well-versed with Pacific culture and heritage and health problems facing the people in the region.

The EU PacTVET project is the first to establish a professional association for resilience practitioners: The Pacific Regional Federation of Resilience Professionals (PRFRP) [23].

This Federation, along with relevant stakeholders like USP, should build capacity regionally to allow the implementation of the SDGs, UNFCCC and the Sendai Framework instruments through grassroots and up to management level. By investing in people’s knowledge and their will to provide safer and more secure environments for their families, friends and visitors alike, it is believed that this intervention may turn the tide for resilience in the Pacific and be able to change how people live and work, thus building more sustainable Pacific communities now and in the future. Clearly, by its very nature, “vocational” education is linked to employment. However, in small Pacific island communities, paid employment is not the norm, therefore, the Resilience qualifications work at the grassroots level by providing training for “productive activities” within the community which will improve resilience, livelihoods, health and well-being. One example of such community activities would be training to ensure the production of safe drinking water – solar water disinfection (SODIS).

SODIS is a process of using solar energy from the sun to destroy the pathogenic microorganisms that cause water-borne disease so that the drinking water treatment can be available at a low-cost solution at the household level [43]. The project was implemented in South Tarawa, Kiribati by the Global Climate Change Alliance: Pacific Small Island States project. SODIS uses readily available resources i.e. sun, 1.5 litres plastic PET bottles and roofing iron to disinfect water, thereby ensuring sustainability of the skills imparted. Communities in South Tarawa were trained in SODIS and awareness activities included the distribution of starter packs and educational games specifically designed for children. SODIS was launched in October 2014 and by February 2015 76% of households in the target communities were using SODIS. The communities reported:

- positive effects including decreases in diarrheal disease especially in children under 5 years old;
- fewer days of school missed;
- decreased spending on kerosene for boiling water;
- better tasting and smelling water compared to boiling water;
- the health clinic servicing the communities also reported decreased incidences of diarrhoea and respiratory illness [43].

SODIS training is needed by most communities in the Pacific who have to deal with lack of safe drinking water on a daily basis.

This can be easily achieved by vocational education in the Pacific because resilience education is a route to better chances in life. Fundamentally, if people chose to pursue better chances in life, then it can be translated into better health outcomes because this has been proven in other areas of life in the Pacific where people have been dealing with cyclones and sea level rise. By training people in the region to be more climate and disaster resilience may not only help them develop innovative solutions to build resilience in health to climate change and disaster caused by natural hazards but also to be revolutionized in turning these threats or impacts into a multipath embedded opportunistic resilience [44, 45].

For example, as a management level, a person with a degree in economics in Tonga may know more about the Tongan economy than health, climate change and hazards for that matter. If the same person enrolled in the course on resilience and health, this person will be exposed to health, climate change and hazards education and will also be able to make an association to economics in order to achieve better livelihoods, health and well-being for his or her family.

### Vocational education is a stepping stone to regionalize and nationalize climate change and health at primary and secondary levels to develop a sustainable solution for the future

If the Pacific is planning to regionalize and nationalize climate change at the vocational education level now and in the future, then this needs to be acknowledged and climate change and health needs to be considered in a different way to exert leadership and take forward global action on addressing climate and health in the region. The best way to do this in the vocational education sector is to use it as a platform as well for the primary and secondary schools to develop a sustainable solution for the Pacific community, thus helping to build resilient Pacific Islanders and communities by 2030 and beyond. From what is known in the region, people’s health suffers because of the impacts of climate change on their health in where they live and work [14] and due to lack of support from the education sector. The end goal of the EU PacTVET project and its PRFRP – and its follow up – is to change this reality.

The task of the EU PacTVET programme is not only to identify and support the application of climate change and health interventions that will do the most to improve the climate change conditions that determine health for the people in the region, but also to help the vocational, primary and secondary education sector progress towards that ideal. These climate change and health adaptation strategies do not just impact on children or young people mortality. Rather they have a powerful impact on adult mortality as well, so powerful that a poor person at the age of 12 in Vanuatu or Tonga has far fewer years in front of him/her than a poor person of the same age in the United State of America (USA) with the same impacts of climate change on their health. In doing so, not only does the education sector change the impact cycle of climate change on health, but it will also help the people of the Pacific to improve their health adaptation strategies in their communities – to restore better health and well-being.

As Queen Salote Tupou III of Tonga stated: “ ‘Oku ‘auha hoku kakai he masiva ‘ilo (the demise of my people is caused by lack of knowledge). This may have been meant for the people of Tonga, but this conceptualisation of knowledge is universal and as a result, it can be applied to other Pacific Islanders as well. The same advice was given by Hosea 4:6 in the Bible that “My people are destroyed for lack of knowledge. Because you have rejected knowledge, I also will reject you from being My priest. Since you have forgotten the law of your God, I also will forget your children” [46].

Importantly, in the Pacific in terms of climate change, health and religion in the region. However, the EU PacTEVT project has responded to this call and will tackle the Pacific lack of knowledge on climate change and health using this new regime. This regime may not only increase the adaptation space for wisdom, skill and knowledge-base for basic education on climate change and health Pacific-wide but will have impacts on other sectors as well (agriculture, coastal management, energy and infrastructure, fishery, forestry, tourism, water resources) since they are interrelated, thus helping vocational, primary and secondary education to develop more sustainable resilience solutions for the future.

## Conclusion

Using vocational education to provide development solutions in the Pacific on health and climate change is new. As a result, there is an opportunity here for the EU PacTVET project to follow through and change this reality once and for all in order to model the degree to which the Pacific society delivers a good life to its citizenry. If the higher education sector already proved it to the world that their strategies worked, then this process should be completed by using the bottom-up approach so that the adaptation strategies process can benefit all levels of society (e.g. streamline climate change and health at primary, secondary and vocational education). In doing so, not only will it landscapes climate change in a way that has never done before, but it will create ownership of the climate change regime and benefit the adaptation space of the Pacific community at large.

There are three ways to achieve this goal. First, is to design a training programme that focuses on training experts in health adaptation through vocational education. This is the best solution that TVET education could provide for climate change and health regarding their participants as students. Second, after designing the training programme to focus on climate change and health, the next phase is to ensure that they get a job. Meaning the programme should be able to provide accredited qualifications on resilience in health to work as climate change advisors or consultants inter alia in their own countries. This is very important for the people in the region because there are two-way benefits: health and economy. Thirdly, when the vocational education is manageable with its programme in situ, the last phase is to help regionalize and nationalize climate change and health at primary and secondary levels for the region to develop more sustainable resilience solutions for the future. Since the Pacific is the most vulnerable region in the world to be affected by climate change and disasters caused by natural hazards, therefore without any doubt, this is the way forward that should be engaged in order to change this reality and reverse this impact of climate change on health.

To ensure that this intervention is achievable for the Pacific, based on the result of the EU PacTVET project, this paper recommends the following policy proposals for the region:

1. to integrate climate change and health into the Pacific vocational education curriculum;
2. to integrate climate change and health into the primary and secondary schools curriculum in the Pacific.

## Acknowledgements

This research was supported by the EU PacTEVT project from the Pacific Community (SPC), The University of the South Pacific, Suva, Fiji and the School of Humanities (Geography), at Bishop Grosseteste University, Lincoln, UK.

